# Competitive Amplification Networks enable molecular pattern recognition with PCR

**DOI:** 10.1101/2023.06.29.546934

**Authors:** John P Goertz, Ruby Sedgwick, Francesca Smith, Myrsini Kaforou, Victoria J Wright, Jethro A. Herberg, Zsofia Kote-Jarai, Ros Eeles, Mike Levin, Ruth Misener, Mark van der Wilk, Molly M Stevens

## Abstract

Gene expression has great potential to be used as a clinical diagnostic tool. However, despite the progress in identifying these gene expression signatures, clinical translation has been hampered by a lack of purpose-built. readily deployable testing platforms. We have developed Competitive Amplification Networks. CANs to enable analysis of an entire gene expression signature in a single PCR reaction. CANs consist of natural and synthetic amplicons that compete for shared primers during amplification, forming a reaction network that leverages the molecular machinery of PCR. These reaction components are tuned such that the final fluorescent signal from the assay is exactly calibrated to the conclusion of a statistical model. In essence, the reaction acts as a biological computer, simultaneously detecting the RNA targets, interpreting their level in the context of the gene expression signature, and aggregating their contributions to the final diagnosis. We illustrate the clinical validity of this technique, demonstrating perfect diagnostic agreement with the gold-standard approach of measuring each gene independently. Crucially, CAN assays are compatible with existing qPCR instruments and workflows. CANs hold the potential to enable rapid deployment and massive scalability of gene expression analysis to clinical laboratories around the world, in highly developed and low-resource J settings alike.

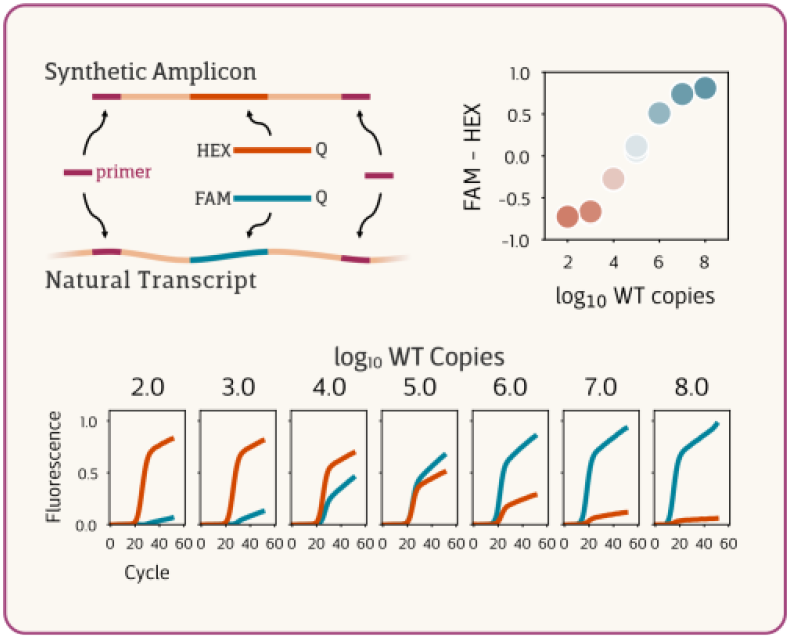

## Introduction

Over the past two decades, gene expression signatures have demonstrated the potential to become a near-universal modality for diagnosis and prognosis of disease. Researchers have discovered a wealth of differential expression patterns that are predictive of a wide range of conditions, from heart disease to infectious disease, tumor chemotherapy sensitivity to stroke recovery prognosis.^1–11^ Despite the progress in identifying these gene expression signatures, however, clinical translation has been hampered by a lack of purpose-built, readily deployable testing platforms. In most cases, signature discovery has been achieved through RNAseq, microarray, or Nanostring analysis, but these instruments require extensive and specialized preparation of the sample, expensive and single purpose instruments, and have yet to find efficient integration with clinical pipelines. Alternatively, some diagnostic gene expression signatures have been implemented by measuring each transcript individually through a set of anywhere from five to fifty parallel qPCR tests, but this approach suffers from low throughput, specialized workflows, high sample volume and reagent use, and poor reproducibility due to the laborious (and often manual) process of dividing a sample across numerous reactions at once. From a clinical perspective, the granularity of these approaches in reporting the exact quantity of every gene in the signature is often unnecessary, as these values are typically meaningless in isolation The only useful information needed is the overall diagnostic indication.

We have developed a simple yet conceptually radical approach to gene expression signature testing that embeds biomarker detection and statistical interpretation directly into the molecular design of a single qPCR reaction **(Figure 1)**. Through careful design of primers, fluorescent probes, and synthetic amplicons, we have engineered molecular tests that detect and interpret the abundance of multiple RNA sequences according to an *in silico*-derived statistical model. For example, we could use synthetic DNA to hard-code the threshold concentrations that separate the “healthy” from “disease” expression patterns for each gene. The reaction then compares the RNA levels in a patient sample to this pre-defined pattern, assessing the degree to which each is more similar to the “healthy” or “disease” populations. Each transcript contributes towards a two-color fluorescent signal, where one color indicates “healthy” and the other “disease”. The relative strength of these two colors depends on the strength of those similarities, calibrated to exactly mimic the components and conclusions of the statistical model.

**Figure 1.**
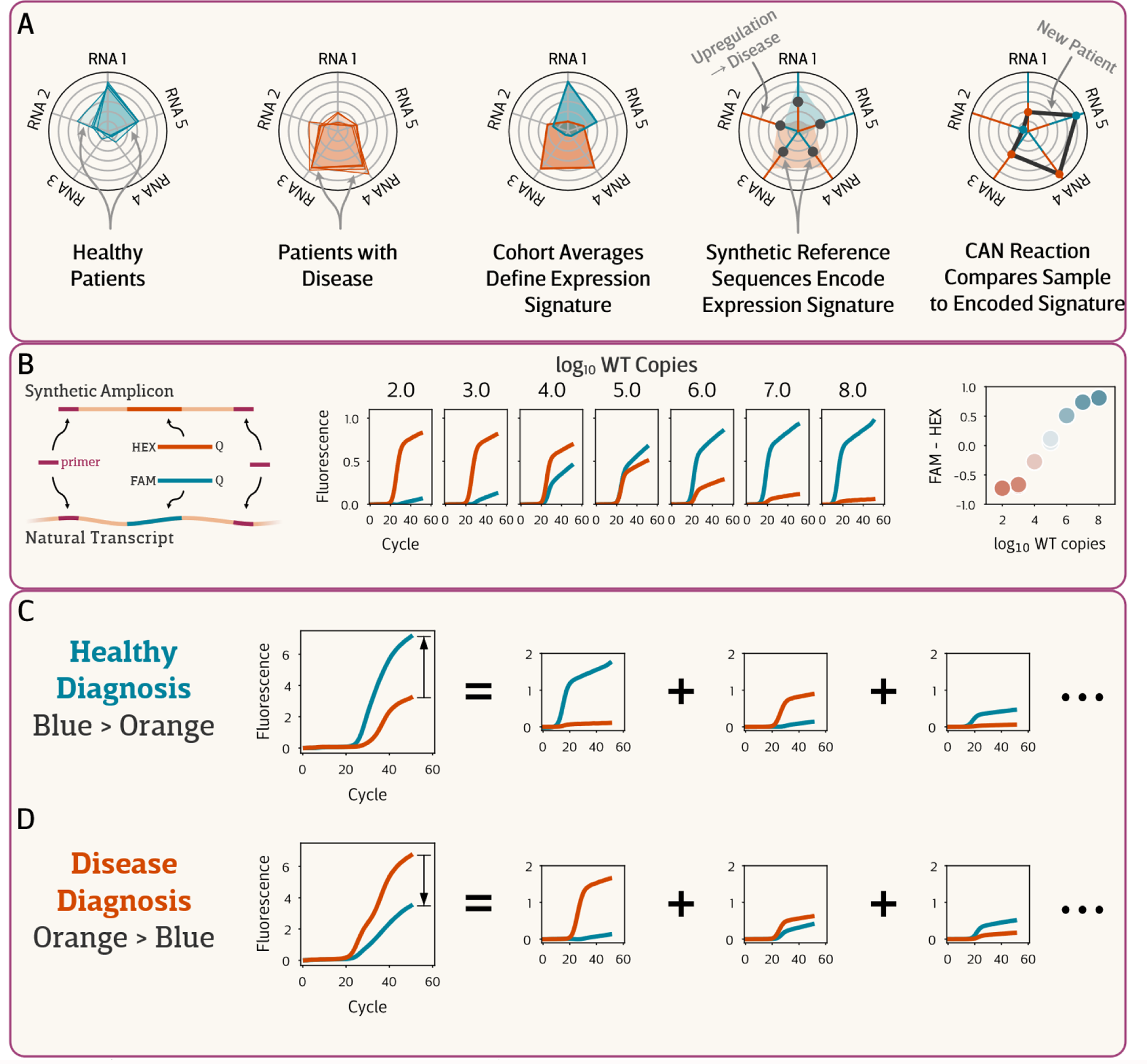
**(A)** Competitive Amplification Networks (CAN) encode and evaluate gene expression signatures. A gene expression signature is composed of RNA transcripts differentially expressed between healthy patients and patients with a given disease or condition. The pattern of expression comprising this signature is encoded within a CAN through inclusion of large synthetic DNA fragments at defined concentrations. The CAN architecture defines not only the per-transcript threshold between disease and healthy populations but also which diagnosis is implied by up- or down-regulation. To assess a new patient, the CAN evaluates each signature RNA in parallel via transcript-specific reaction modules; the further away a given transcript is from its threshold, the greater the evidence for that diagnosis and the stronger the signal produced. The signals from all of these parallel comparisons combine additively, and diagnosis is achieved by whether more evidence (a stronger signal) points towards disease or healthy. **(B)** An individual CAN reaction module consists of a natural transcript, one or more synthetic amplicons, distinct probes labeled with one of two colors, and primers common to both natural and synthetic targets. At a fixed concentration of the synthetic amplicon, PCR amplification of different transcript concentrations leads to modulation of the steady state intensity of both probe fluorophores. The difference between these two intensities provides a signal that indicates whether the transcript is above or below a specified threshold. **(C, D)** In the full CAN, the steady-state fluorescent intensities from all individual modules combine additively. The CAN is designed and calibrated such that diagnosis corresponds to whichever fluorescent signal is more intense.

These novel PCR architectures, which we refer to as Competitive Amplification Networks (CANs), act as biological computers,^12^integrating evidence to perform Bayesian inference on a molecular level and arrive at a diagnostic conclusion. The key advantage of this one-pot reaction is its compatibility with commercial reagents, ubiquitous qPCR thermocyclers, and standard clinical workflows, allowing these tests to be easily deployed to diagnostic labs in highly developed and low-resource settings alike.

Here we describe the design principles for these Competitive Amplification Networks. We first demonstrate how competitive PCR can be engineered to produce a tunable fluorescent signal proportional to the relative initial abundance of the two amplicons. To tackle the richness of this design space, we describe our mathematical framework for characterizing amplification behavior, mechanistically predicting the ideal design of CAN components towards a given task, and statistically optimizing competitive reaction modules. Next, we demonstrate how higher-order CAN architectures enable additional functionality and flexibility in the signal response profile. We illustrate how numerous reaction modules can be combined in a single reaction through development of CAN systems for each of a two- and a four-gene expression signature. Finally, we evaluate the clinical performance of a diagnostic CAN test for distinguishing fevers of bacterial and viral origin based on patient blood samples, demonstrating perfect agreement with the gold standard approach.

## Results

Our Competitive Amplification Networks are organized around modular reaction components specific to each target sequence. Each module in a CAN is built on the principles of competitive PCR,^13-15^ where multiple distinct amplicons compete for binding to the same primer(s). Here, we design synthetic, exogenous DNA sequences to bear the same primer-binding sequence as the natural, endogenous RNA targets **(Figure 1B)**. Previously, endpoint competitive PCR was used as an alternative to “real-time” fluorescent PCR for target quantitation; we fuse the two approaches by designing distinct fluorescent probes for both natural and synthetic amplicons. The difference in endpoint signal intensity between these two colors constitutes the reaction readout.

These design principles create the possibility of multiple reaction module configurations that differ in the number of natural and synthetic targets and the arrangement of primers shared between them. We outline several reaction module architectures below, demonstrating the distinct behavior and design landscape of each, before illustrating how multiple modules can be combined to produce a complete assay.

### Model-driven module engineering

To explore the rich design space of CAN systems and support the optimization of specific implementations, we developed a framework of mechanistic and statistical modelling. Competitive Amplification Networks rely on the interactions of multiple replicating sequences competing for limited pools of resources, namely primers. The key to predicting the behavior of this complex system is a mathematical model that can translate observations of each sequence amplifying in isolation to the rich dynamics that arise when they interact with each other. Existing models of PCR are either too simple to allow prediction of interactions or too complex to allow complete parameter estimation from individual observations.

We introduce the use of Monod-like growth kinetics for characterizing PCR dynamics. The classical Monod model describes how bacterial growth is limited by resource consumption via a system of ordinary differential equations. Theoretically, every target strand in a PCR reaction is duplicated with each thermal cycle, yet this clearly won’t hold true for very long sequences nor as the amplicon concentration approaches the remaining primer concentration. We adapt Monod kinetics to describe how the amplification rate of a given target sequence evolves during the reaction as key substrates (in particular primers) are consumed (full model details are given in the **Supplementary Information)**. Our kinetic model successfully recapitulated observed reaction dynamics **(Figure 2A)**, allowing *in silico* exploration of various CAN configurations.

**Figure 2.**
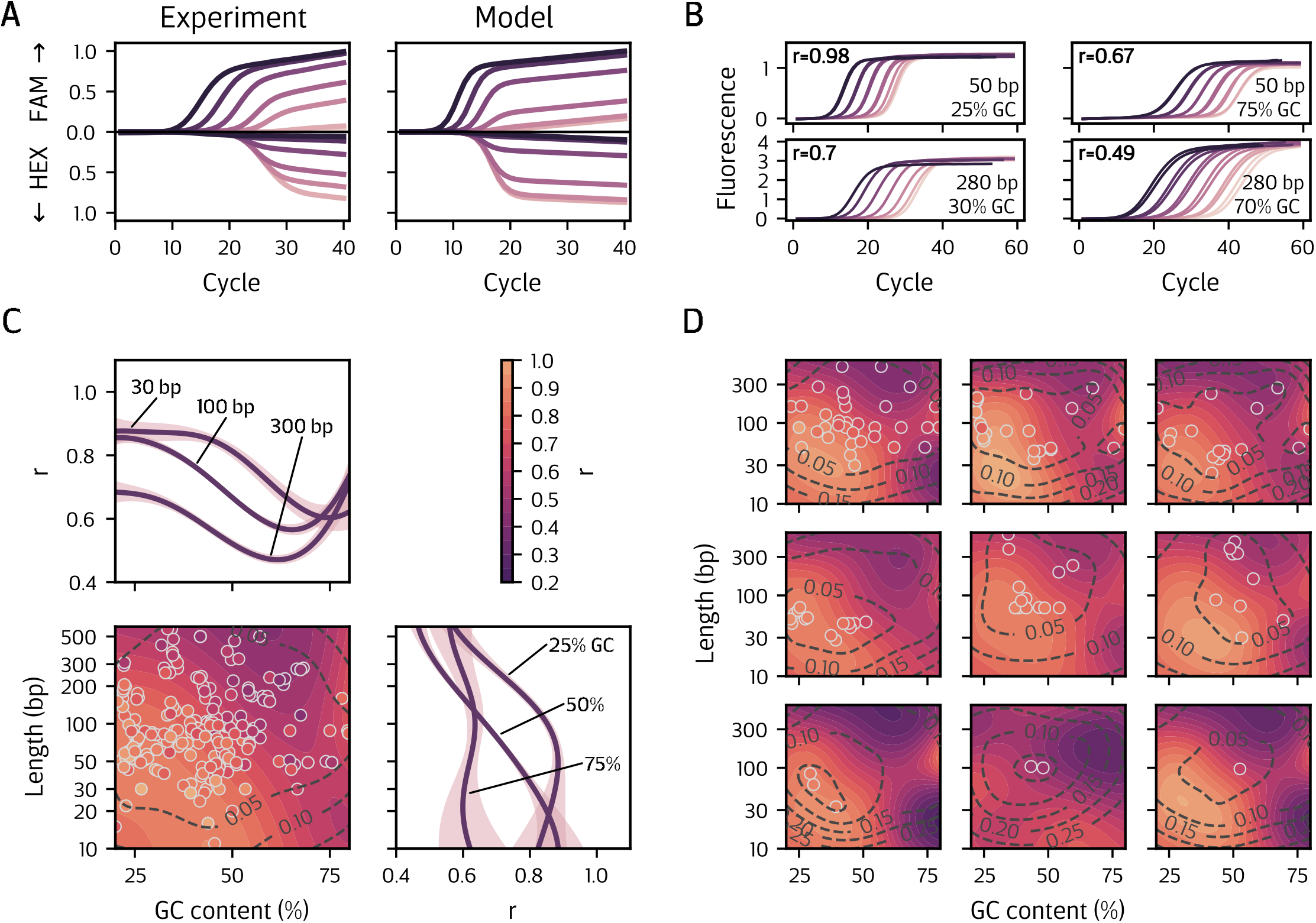
**(A)** Our kinetic model captures competitive dynamics observed experimentally. This model allows us to parameterize the amplification behavior of individual sequences and provides the basis for simulating various competitive configurations. **(B)** One key model parameter, amplification rate r, influences the “steepness” of the amplification curve. Shown here are four different amplicons that vary in length and GC content amplified with the EvaGreen dye. Because EvaGreen fluorescence results from intercalation into the double helix, the final fluorescence intensity is strongly correlated with the length of the amplicon. For probe-based detection, the same association is not observed (not shown). **(C)** Amplification rate depends strongly on the length and GC content of the amplicon. Here we fit a Gaussian Process (GP) to the observed rate of 305 tested sequences. The GP captures global trends along with uncertainty resulting from both biological variation/noise and data sparsity. The colored surface reflects the GP mean while dashed lines denote contours of the GB standard deviation. The plots above and to the right show slices of the regression surface (median with 95% Cl) at the indicated positions. **(D)** We also fit a GP with the Linear Model of Coregionalization, which treats the observed sequences as belonging to distinct, but correlated, surfaces defined by a given primer pair. Each surface shown here reflects multiple sequences designed with a given primer pair. This approach can better capture the nuances of the relationship between amplicon length, GC content, and rate that are specific to a given primer pair while allowing those surfaces to share information about the global trend.

To design natural and synthetic sequences to exhibit specific amplification behaviors, we developed a statistical model based on Gaussian Process (GP) regression.^16^ GPs have shown promise in modeling and engineering biological systems previously.^11-20^ Our simplest GP model relates the length and guanine-cytosine (GC) content of all sequences tested to the kinetic parameters fitted to their amplification profiles, in particular the rate *r* **(Figure 2C)**. However, this “global” model has the disadvantage of obscuring the outsized impact of the primer regions (underfitting). Considering only targets designed with a given pair of primers resulted in systematic deviations from the global trend, yet this approach ignores commonalities in behavior between primer pairs, resulting in overfit surfaces **Figure S1**. To achieve a middle ground between these two extremes, we augmented our GP with a Linear Model of Coregionalization **(Figure 2D)**.^**21**^ This technique assigns a distinct regression surface to each primer pair yet learns the correlation between observations from different surfaces. This allows primer pairs with many different tested sequences to inform the GP’s predictions of primer pairs with only one or a few sequence observations. Crucially, this “partial pooling” approach exhibits well-calibrated uncertainty, so while a general trend is assumed to hold true for surfaces with few observations, low confidence is assigned to regions far from the actual observations.

New sequence designs were selected through the Expected Improvement (El) algorithm. The algorithm is given an objective value and a probability surface learned through regression, for instance one that relates sequence length and GC content to observed amplification rate, From this surface, EI selects a new point for investigation (length and GC content of a novel amplicon sequence) by considering both the magnitude of its anticipated improvement over the previous observations towards the objective value and the probability of achieving that improvement Our full design-test -build workflow was as follows. **(1)** Use kinetic simulations to identify the ideal amplification behaviors for a given task. (2) Observe the amplification of a set of sequences. (3) Estimate kinetic parameters for these observations. (4) Relate heuristic descriptions of these sequences (primers, length, GC content) to their measured parameters with a GP. (5) Use EI to select the sequence heuristics most likely to achieve the desired behavior. (6) Use sequence design tools (Nupack^22 23^) to design synthetic PCR targets with a given length, GC content, and other sequence constraints. By integrating both theoretical and empirical modeling, this workflow proved very amenable to biological optimization in a low-data setting.

### Bipartite Module

The simplest CAN module consists of a single synthetic amplicon that shares both primer regions with a single target sequence **(Figure 3A)**. When the natural and synthetic targets are amplified in the same PCR reaction, they compete for the same primers. Since primers are consumed by each replication of a target strand, the amplification of both sequences stops as soon as the primer pool is exhausted. The quantity of each amplification product at the end of the reaction thus depends on the relative starting quantity of the two targets.

**Figure 3.**
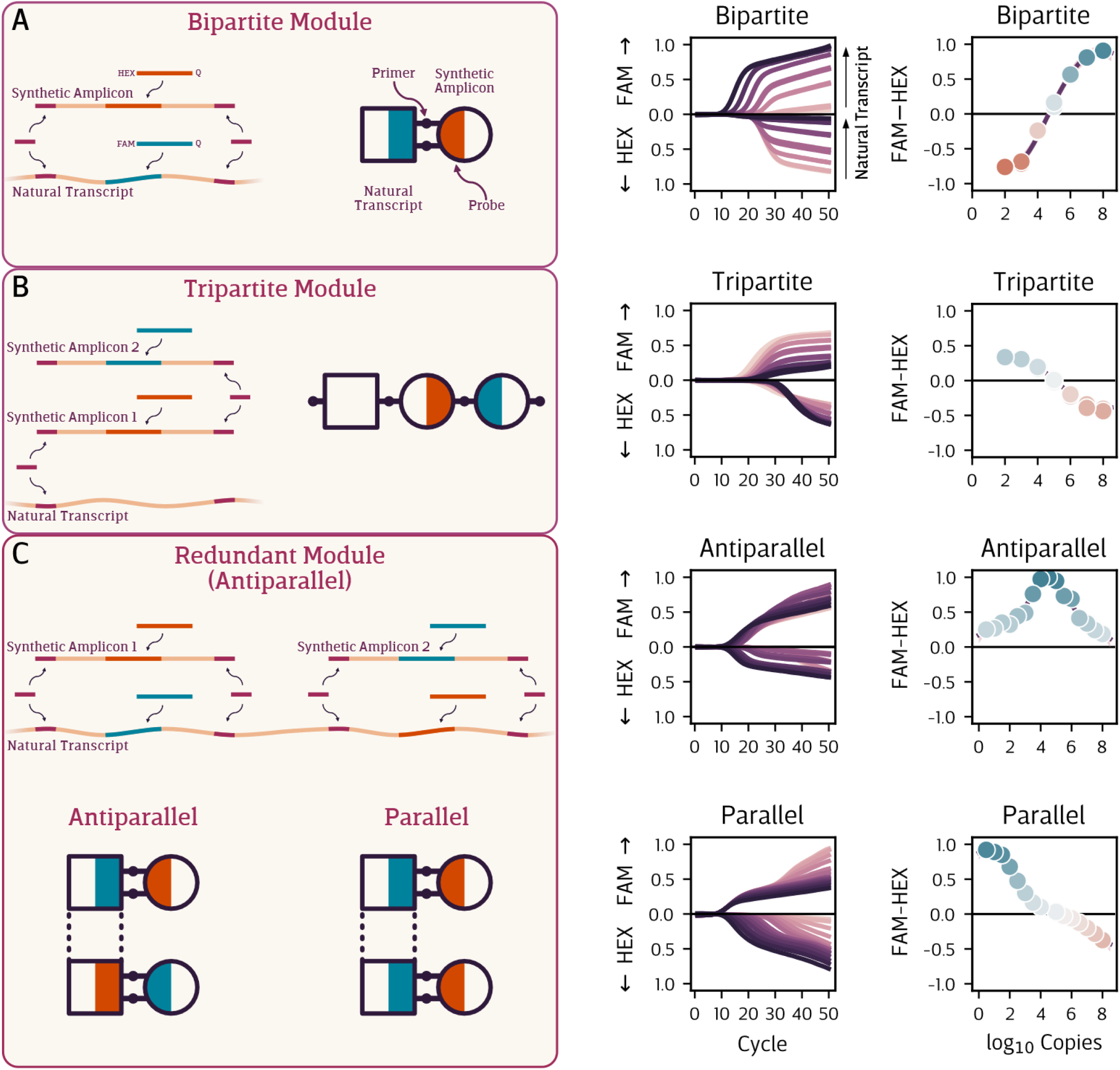
Competitive Amplification Networks consist of a collection of reaction modules, each one tailored to a given RNA transcript, miRNA sequence, or other nucleic acid target. These modules leverage competition between natural and synthetic amplicons for the same pool of primers (left column), resulting in a two-color fluorescent signal (middle column) that reflects the initial concentration of the natural target (right column). **(A)** The “bipartite” module uses a single synthetic competitor to generate a locally-linear response profile. **(B)** The “tripartite” module uses a second synthetic competitor, producing a similar profile to the bipartite module but without the need to design a probe for the natural sequence. **(C)** “Redundant” modules target a single transcript in multiple locations, allowing highly non-linear, non-monotonic, multi-phasic, and multi-modal response profiles. The design flexibility of these modules enables nearly arbitrary response profiles.

The synthetic target is designed to differ from the natural target in the “core” region between primers, allowing distinct fluorescent probe oligos to be designed specific to each. These probes translate the difference in amplicon abundance at the end of the reaction to a difference in intensity between the respective fluorophores. At a fixed concentration of the synthetic amplicon, the magnitude of this difference follows a sigmoidal relationship with the initial concentration of the natural target. For example, if a FAM probe is used for the natural target (which can be designated *NF)* and a HEX probe for the synthetic target (designated *A*^*H*^), the HEX signal dominates when there is less natural target than synthetic at the beginning of the reaction ([*N*^*F*^]_o_ « [A^*H*^]_o_) and the FAM signal dominates when the opposite is true ([*N*^*F*^]_o_ « [*A*^*H*^]_o_) Between these two extremes ([*N*^*F*^]_o_ “[*A*^*H*^]_o_), a smooth transition from primarily HEX to primarily FAM signals occurs, providing an approximately linear signal regime.

The signal response of a bipartite CAN module can be tuned through various design choices **(Figure S2)**. By assigning a FAM probe to the natural target and a HEX probe to the synthetic competitor (*N*^*F*^:*A*^*H*^), the FAM-HEX signal will increase with the concentration of the natural target; swapping the fluorophores *(N*^*H*^: *A*^*F*^) will give a signal that decreases as the concentration of the natural target is increased. The concentration of primers and probes determines the magnitude of the response, with lower amounts leading to a weaker signal. The concentration of the synthetic amplicon affects the midpoint of the signal, at which the two fluorescent colors are equal in intensity. Finally, the exact sequence composition of the synthetic and natural amplicons (and, to a lesser extent, the primers) influences both the sharpness of the sigmoidal response and the signal midpoint.

### Tripartite module

A key limitation of the bipartite module is the need for a distinct probe to be designed for every new natural target. The tripartite module overcomes this constraint through inclusion of an additional synthetic sequence **(Figure 3B)**. Here, the natural target *N°* has no probe and competes with a HEX probe synthetic target *A*^*H*^ for only one primer; the second primer specific to the natural target is “uncontested”. A FAM-probe synthetic target *B*^*F*^ is also included, designed to compete for the remaining primer from *A*^*H*^; the remaining primer for *B*^*F*^ is also left uncontested. In all, the tripartite module consists of three amplicons (one natural, two synthetic), four primers (two contested and two uncontested), and two probes.

In this *N*^*°*^*:A*^*H*^*:B*^*F*^ configuration, the observed signal comes only from the synthetic amplicons, yet for a given concentration of *A*^*H*^ and *B*^*F*^ the response profile is still dependent on the initial concentration of the natural target **(Figure S3)**. In this way, the natural target can be seen as simply biasing the amplification equilibrium of the two synthetic targets. As the concentration of *N*^*°*^ is increased, *N*^*°*^ applies additional competitive pressure to *A*^*H*^, allowing *B*^*F*^ to eventually outcompete *A*^*H*^ and causing the FAM signal to dominate.

Achieving an informative response profile with a tripartite module required careful design. Computational optimization of the kinetic model for this CAN architecture suggested the natural target *N°* needed to exhibit an amplification rate *r* considerably greater than the synthetic competitors *A*^*H*^ and *B*^*F*^ (specifically *r*_*N*_ *= 0*.*8* and *r*_1_=*r*_2_=0.6). Indeed, an implementation of this module with *r*_*N*_*=0*.*82, r*_1_=0.52, and *r*_2_=0.5 produced the desired signal response **(Figure S4)**, while an implementation where all three rates were similar (*r*_*N*_=0.71, *r*_*1*_*=0*.*64*, and *r*_2_=0.65), led to minimal signal change across *N°* concentrations **(Figure S4)**.

### Redundant competition

The bipartite and tripartite competitive modules outlined above are limited to monophasic, monotonic signal response profiles: they exhibit approximately linear transitions from their minimum to their maximum signals. However, more complex behavior can be achieved by designing multiple independent submodules for different regions of a single transcript producing a “redundant” composite module **(Figure 3C)**.

The “parallel” configuration of the redundant module consists of two bipartite submodules where both natural amplicons are designed with FAM probes and both synthetic amplicons are designed with HEX probes (*N*_*A*_^*F/*^:*A*^*H*^+*N*_*B*_^*F*^:*B*^*H/*^) .Varying the concentrations of the synthetic amplicons produced response profiles with two distinct transition regimes, differing in either sharpness or position relative to the concentration of the natural target **(Figure S5)**. In the “antiparallel” configuration. one submodule used a FAM probe for the natural amplicon and a HEX probe for the synthetic. while the other submodule was designed the opposite way (*N*_*A*_^*F/*^:*A*^*H*^+*N*_*B*_^*H*^:*B*^*F*^). This configuration produces a signal which tends in one direction then reverses course, producing a peaked response profile reminiscent of a band-pass filter. The concentrations of the synthetic amplicons determine the direction, breadth, and position of the peak.

### Combining Modules

Full implementation of a CAN consists of integrating multiple modules together in a single reaction mixture. When optimized appropriately, each module produces an independent contribution to the overall signal **(Figure 1C, D)**. The result is a multidimensional response surface, where each combination of target concentrations produces a characteristic (though non-unique) signal.

In the gene expression signature setting, each target is a gene transcript whose abundance is known to correlate with disease state **(Figure 1A**). Often, this statistical relationship is derived from logistic regression,^24^ which finds a linear relationship between expression level of each gene in the signature and its contribution to the log-odds of one category over the other.^25^ To mimic this statistical relationship, a CAN reaction module can be designed for each transcript. Each module produces a two-color fluorescent signal that is approximately linear with respect to target concentration within a certain window. By optimizing the module components, this fluorescent signal can directly reproduce the linear relationship derived from logistic regression. Finally, combining multiple modules in the same reaction produces a composite two-color signal where each color represents the additive contributions from each individual module. This is fundamentally the same principle as logistic regression, where the positive or negative contributions of each predictor are added together to give the ultimate log-odds of the respective disease states. The CAN system integrates evidence provided by each transcript to present not only which disease state is more likely but also the *strength* of that conclusion. In essence. CANs perform Bayesian inference^26^ on a molecular level.

To illustrate this concept, we designed a CAN comprised of four bipartite modules to reproduce a four-gene expression signature for diagnosis of tuberculosis.^2^,^27^ We tested this system over an approximately 5x5x5x3 grid of target concentrations **(Figure 4)**. The resulting signal was dependent on the concentrations of all four targets. Nonlinear regression revealed that the “marginal” contribution from each target was largely linear and sloped, suggesting each module retained its independent behavior when incorporated into the larger CAN. Thus, this system can accurately reproduce a statistical linear model with four predictors.

**Figure 4.**
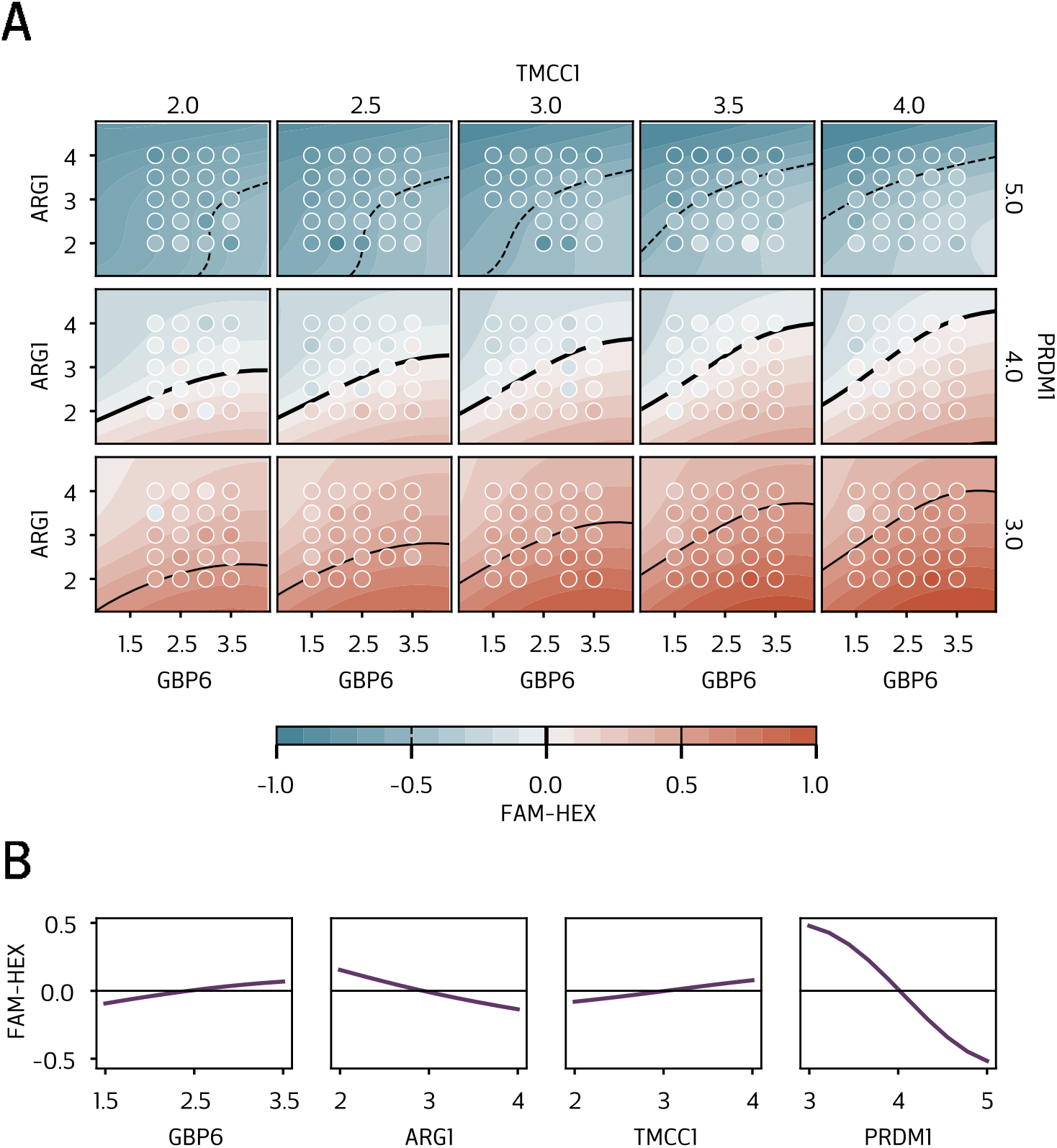
Multiple reaction modules can be combined, resulting in an multidimensional response surface that integrates the information from all targets. **(A)** Here we combined four different “direct” modules, one each for the natural targets GBP6, ARGl, TMCCl, and PRDMl. We tested this reaction mixture against various combinations of the four targets in an approximately 5x5x5x3 grid (respectively). Each filled circle is the total FAM HEX response for a single reaction. The smooth surface is the mean of a GP fit to all reactions: the thin dashed, heavy solid, and thin solid lines represent -0.5, 0.0, and +0.5 signal response, respectively. **(B)** The marginal response to each component, i.e. the average across all other components. The near-linearity of each marginal component verifies that the four-dimensional response is the sum of linear one-dimensional responses. Thus, this system can accurately reproduce a statistical linear model with four predictors.

### Applications

#### Mutant allele detection at low-VAF

A promising application of tripartite CAN modules is in detection of low concentrations of mutant alleles (especially single-nucleotide variants, SNVs) in the presence of a large excess of wild-type alleles. By identifying tumor-derived DNA fragments in the bloodstream (circulating tumor DNA, ctDNA), such low-VAF (variable allele frequency) SNV detection has the potential to aid in early cancer diagnosis or minimal residual disease (MRD) testing for monitoring of cancer treatment efficacy.^28^ Sequencing approaches typically exhibit high costs and turn-around times, dangerously delaying treatment decisions, while traditional PCR approaches lack the capacity to detect the numerous mutant alleles necessary for reliable MRD.^29^

To demonstrate the potential for MRD detection on our platform, we sought to modulate the detection system such that a mutant allele could be detected even when the corresponding WT was present at orders of magnitude greater concentration. This was achieved through incorporation of a non extensible blocker oligo, adapting an approach from Blocker Displacement Amplification (BDA) **(Extended Data Figure 1)**.^30^ This short strand of DNA was designed to be complementary to the WT and partially overlap with the 3’ region of the gene-specific reverse primer. The blocker was synthesized with a C3 modification at the 3’ end, to prevent extension by the Taq polymerase, and a locked nucleic acid (LNA) base at the SNV site, to increase the strength of its binding to the WT allele. During amplification, the blocker competes thermodynamically with the reverse primer: the blocker outcompetes the primer for binding to the WT yet the primer largely outcompetes the blocker for binding to the SNV. Thus amplification of the WT is suppressed while the SNV is amplified relatively unimpeded.

At high blocker concentration, we observed that the SNV produced a signal 10,000 fold greater than the WT: 10^5^-10^6^ copies of WT were needed to significantly alter the signal from 10-100 copies of SNV. Our results demonstrate the potential of our technology to detect mutant alleles at countable concentrations down to 0.1% VAF and up to 10^5^ copies of WT.

#### A two-RNA signature for distinguishing viral and bacterial fevers in infants

To illustrate the clinical utility of the CAN platform, we designed a reaction to reproduce a two transcript gene expression signature that accurately determines whether a fever in an infant is viral or bacterial in origin **(Figure 5)**.^31^ We first performed logistic regression on transcript counts for each gene resulting from Nanostring analysis of 42 patient samples. Logistic regression determined the linear relationship between the abundance of each transcript and its marginal contribution to the “log-odds”, or relative probability, of the two diagnoses.

**Figure 5.**
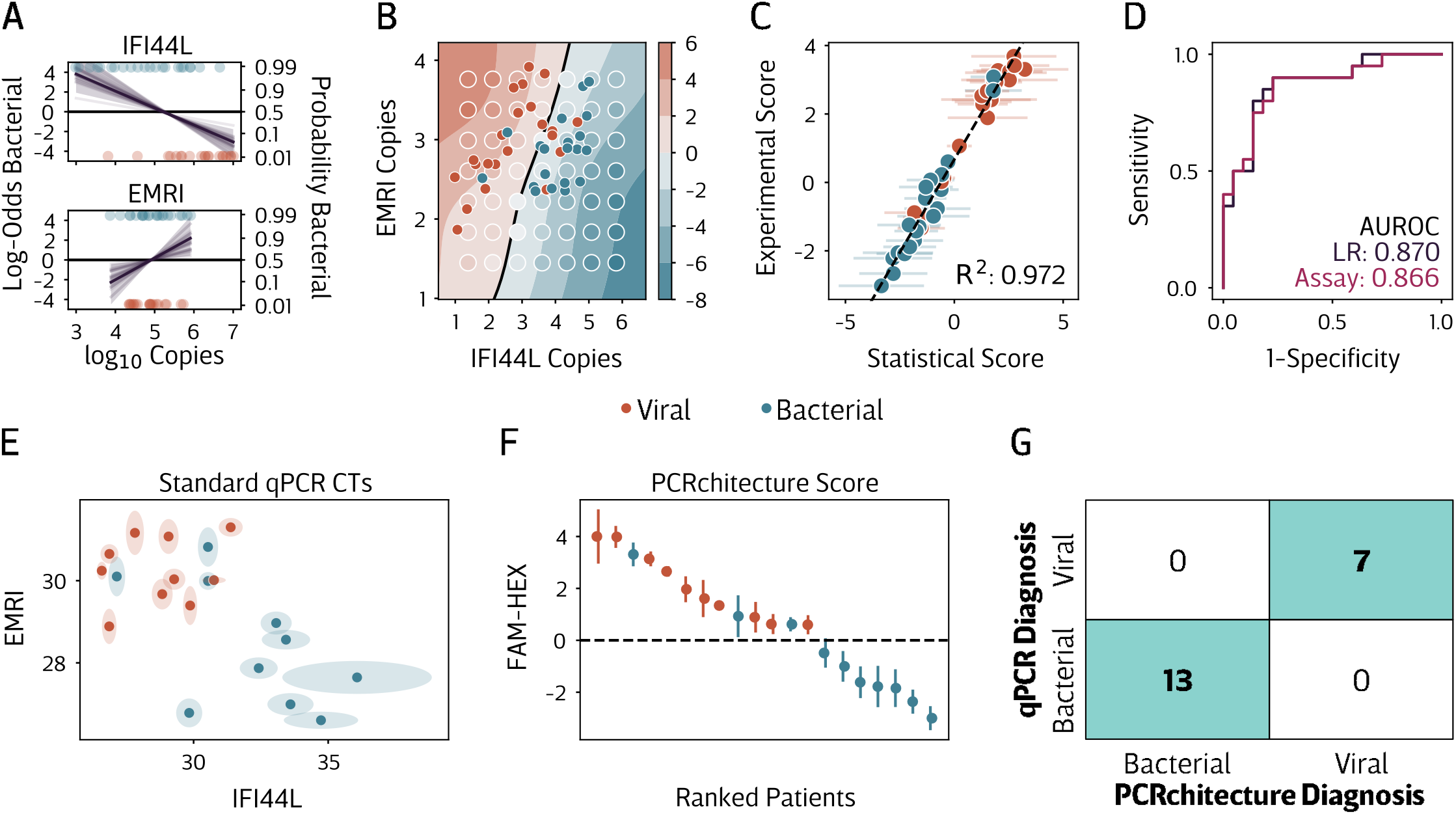
A CAN design reproduces a two-transcript signature for determining fever etiology in infants. **(A-D)** Assay development with synthetic samples. **(A)** Logistic regression of patient data determines the contribution of each transcript to the diagnosis, in this case whether the fever is bacterial or viral in origin. These marginal effects provide the design objective for the reaction modules. **(B)** After optimizing the respective reaction modules, the combined signal is representative of the concentrations of the two target RNAs. Gaussian Process regression (shaded surface) on experimental points (large circles) allowed us to estimate the experimental score for each patient (small dots) in our training set. The decision boundary (black line) at a score of O separates the bacterial (red) from viral (blue) patients. **(C)** This experimental score is tightly correlated to the statistical score resulting from logistic regression, **(D)** exhibiting nearly identical area under the receiver-operator curve (AUROC). These comparisons confirm that the CAN reaction faithfully reproduces the gene expression signature. **(E-G)** Assay validation using clinical samples. **(E)** Performing separate qPCR on clinical samples for each transcript in the signature (shaded area 95% Cl from five replicates) resulted in clustering of patients by diagnosis, albeit with high CT variability and three misdiagnosed bacterial patients. **(F)** Performing the single CAN assay on the same patients achieved identical separation by diagnosis (means w/ 95% Cl from five replicates) yet in a single reaction and with lower variability than qPCR CTs. **(G)** Per patient diagnosis was identical between approaches.

We then designed bipartite competitive reaction modules for each of the two transcripts, tuned such that their individual response profiles recapitulated the marginal contributions identified by logistic regression. Finally, we tested this preliminary assay design on 20 patient samples and compared against the “classical” PCR approach of separately quantifying each RNA target **(Figure SE-G)**. Our CAN-based assay produced identical diagnostic conclusions for every patient. Furthermore, our assay exhibited improved reliability over the classical approach.

This can be attributed to the naturally self-limiting nature of competitive amplification: the sigmoidal response profiles limit the influence of outliers, while classical C_T_s can vary considerably at low concentrations. These results demonstrate the ability of the CAN system to condense the power of gene expression signatures into a single PCR reaction with immediate clinical utility.

## Discussion

To achieve diagnosis and prognosis for complex diseases, we need to develop molecular tests capable of integrating information from multiple biomarkers simultaneously. When faced with multiple biomarkers, as in gene expression signature diagnostics, the traditional analytical approach has been to simply measure each individually, a paradigm that scales poorly. This approach relies on either expensive, specialized equipment that restricts analysis to well-funded, centralized hubs or laborious parallel testing methods that are difficult to implement, exhibit poor reproducibility, and create analysis bottlenecks. Philosophically, individually measuring each target is unnecessary; it ignores the fact that the only relevant information is the pattern as whole.

We developed Competitive Amplification Networks to enable such pattern recognition on a molecular level. CANs are designed as a set of target-specific modules, each of which evaluates the expression level of its target in the context of known “healthy” and “disease” patterns. To permit a range of different design constraints, we developed and characterized a suite of CAN module archetypes. In the bipartite module, a single synthetic amplicon is designed to share both primers with the natural target of interest. Including distinct fluorescent probes targeting the synthetic and natural amplicons produces a sigmoidal response profile over different concentrations of the natural sequence. The approximately linear region in the middle of this response can be used to recapitulate a target specific contribution in a linear statistical model. In a tripartite module, a second synthetic amplicon is included, while the first is designed to share one primer with the natural target and its remaining primer with the other synthetic amplicon. This module has the advantage of needing probes only for the synthetic amplicons. This not only expands the design space available but could reduce development cost for a large signature considerably, as the entire network could theoretically be designed with only four probes in total, independent of the number of natural targets. Finally, redundant competitive modules comprise several submodules specific to distinct regions of a single target sequence. This design strategy enables engineering of more complex response profiles, such as multi-phasic signals and “bandpass filters”.

Multiple CAN modules can be combined to produce a multi-dimensional response that integrates the component one-dimensional responses. We demonstrated the clinical utility of the CAN system by designing an architecture specific to a previously reported gene expression signature for discriminating fevers of viral and bacterial origin. The experimental results of this CAN demonstrated perfect agreement with a gold standard approach which measured each transcript independently. These results support the clinical potential of CANs for allowing gene expression signature diagnosis through a single PCR reaction.

Our approach takes inspiration from developments in synthetic biology and DNA strand displacement, both of which have presented ingenious designs for detection of complex biological signals and patterns.^32-40^ However, these systems have retained two key barriers to implementation as clinical diagnostic platforms. First, they are often slow, requiring several hours to reach steady state. Second, they rely on bespoke reaction conditions that have yet to be mass manufactured and distributed. Our CANs, by contrast, produce an interpretable signal in less than hour. In addition, CANs rely only on commercial TaqMan PCR master mixes, which have already been highly optimized for mass production and worldwide distribution. Manufacture of a disease-specific CAN requires only a mixture of primers, probes, and synthetic amplicons and is thus well-suited to rapid clinical translation.

The flexibility of CAN architectures allows the engineer to embed a statistical model into the network design. The most straightforward implementation of CANs employs the mostly linear region of the bipartite or tripartite signal response profile to reproduce a (generalized) linear model, such as the logistic regression model described above. This could likely be extended to other linear and non-linear frameworks.

All the CAN architectures employed here were additive, with each target producing a signal independent of the others. However, the natural evolution of this approach would be to design architectures where target signals are interdependent. In such a design, synthetic amplicons could be used to “link” targets and modulate the signal. This could lead to the engineering of Boolean-like networks that, for instance, produce a positive signal only when two targets are both above some threshold, and a negative signal otherwise. Such higher-order competitive architectures could prove capable of very nuanced recognition of complex biological network states.

## Conclusion

Our CAN system provides the building blocks for capturing a snapshot of an entire biological network state at once, combining detection and interpretation into a single step. We exploit a fundamental principle of biological signaling - competition for limited resources - to integrate and classify an entire pattern of nucleic acid targets. CANs achieve such computation in a simple, one-pot reaction seamlessly compatible with existing clinical instrumentation and workflows. Indeed, our implementation of a CAN for assessment of a gene expression signature exhibited perfect agreement with the gold standard approach on clinical samples. By enabling precision diagnostics around the world, CANs have the potential to dramatically expand access to personalized medicine.

## Materials and Methods

### qPCR conditions

All qPCR reactions were performed in an Applied Biosystems QuantStudio 6 Flex using Applied Biosystems MicroAmp EnduraPlate Optical 384-well plates (Thermo Fisher Scientific, Waltham, MA, USA). Reactions were performed at 10 µL. Thermocycling stages included a 3s melt step at 95 °C and a 30s annealing/extension step at 60 °C. Applied Biosystems TaqMan Fast Advanced Master Mix was used for reactions containing only DNA templates while Applied Biosystems TaqMan Fast Virus I-Step Master Mix. A typical reaction contained 100 nM of each primer and either 150 nM fluorescent probe or lx EvaGreen dye (Biotium, Fremont, CA, USA).

### DNA and RNA

Primers and probes were purchased from Integrated DNA Technologies (“IDT”, Coralville, IA, USA). Competitor sequences and synthetic natural-target analogs were purchased either as eBlock Gene Fragments from IDT or as Gene Fragments from Twist Biosciences (San Francisco, CA). Synthetic RNA was generated through *in vitro* transcription of gene fragments using HiScribe T7 High Yield RNA Synthesis Kit (New England Biolabs, Ipswich, MA, USA) and quantified via qPCR standard curve.

### Sample Collection

Patients were recruited with informed, written parental consent as part of the IRIS (UK) and GENDRES (Spain) studies (ethical permissions: St Mary’s Research Ethics Committee, UK REC 09/H0712/58: Ethical Committee of Clinical Investigation of Galicia (CEIC ref 2010/015). Patients were phenotyped according to their likelihood of bacterial or viral illness according to an agreed algorithm.^31^ Patients were classified as bacterial only if they had a pathogenic organism isolated at a usually-sterile body site. Viral patients had an identified pathogen, a consistent syndrome, and low inflammatory markers (CRP<60mg/L).

### Nanostring Analysis

RNA was extracted from whole blood PAXgene tubes using Qiagen PAXgene RNA extraction kits. Gene abundance quantification was undertaken using the NanoString nCounter MAX system. Raw counts were normalized and log transformed.

### Data Analysis and Modeling

All analysis was performed in the Python programming language, version 3.9. Notable packages used include PyMC3^41^ for Bayesian parameter estimation and Gaussian Process regression: JAX^42^ for differential equation simulations: Numpy^43^, Scipy^4^4, and Pandas^45^ for general data processing: and Seaborn^46^ and Matplotlib^47^ for visualization.

## Acknowledgements

Research reported in this publication was supported by the National Institute Of General Medical Sciences of the National Institutes of Health under Award Number F32GMI31594.lhe UKRI CDT In Al for Healthcare (EP/SO23283/1),. the BASF / Royal Academy of Engineering Research Chair In Data-Driven Optimisation, the EPSRC IRC Next Steps Plus grant (EP/RO187O7/1), the EPSRC Centre for Doctoral Training in BioDesign Engineering IP SO22856 11, and the Royal Academy of Engineering Chair In Emerging Technologies award (CiET2O21\94) Tor the purpose of open access, the author has applied a Creative Commons Attribution ICC BY) license 10 any Author Accepted Manuscript version arising. The content Is solely the responsibility of the authors and does not necessarily represent the official views of the National Institutes of Health or any other funding agency.

## Data and Code Availability

Experimental data and analysis code can be provided upon reasonable request. Requests for patient data must demonstrate a pressing need and adherence to responsible health data stewardship

## Supplement

### A Monod-like model for PCR dynamics

Our Competitive Amplification Networks rely on the interactions of multiple replicating sequences competing for limited pools of resources, namely primers. The key to predicting the behavior of this complex system is a mathematical model that can translate observations of each sequence amplifying in isolation to the rich dynamics that arise when they interact with each other. Existing models of PCR are either too simple to allow prediction of interactions or too complex to allow complete parameter estimation from individual observations.

We introduce a novel model for characterizing PCR dynamics. This model, based on the Monad model of bacterial growth,^32,48^ .has two key advantages over previous models: simplicity and modularity. The most common parameterization of the PCR curve is likely the four- or five-parameter sigmoid,^49^modeled after logistic growth.^24^ While convenient, this model is not modular: it offers no insight into how two amplification systems would interact. In contrast to these previous models, our Monad-like model is simple enough to allow full estimation of all parameters from a single reaction trace yet flexible enough to allow prediction of interacting behaviors.

Our model is constructed from a mechanistic view of PCR amplification yet abstracts away many fine-grained biophysical details. Theoretically, every target strand in a PCR reaction is duplicated with each thermal cycle. However, it is clear that, even under extreme primer excess, the proportion of targets duplicated could be less than perfect. A very long amplicon under a relatively short PCR extension phase, for example, would likely fail to amplify efficiently. Whatever this maximum proportion is, it will not hold true for the entire reaction: as the concentration of target strands approaches and exceeds that of the remaining primers, the proportion of targets duplicated will naturally decrease. Other amplification resources, such as oligonucleotide probes or nucleotide triphosphates, could also limit target replication. Notably, in the competitive or multiplexed PCR setting, various resource pools will be consumed by multiple different targets simultaneously. We chose the Monad model of bacterial growth as a basis for our PCR model due to its ability to capture all of these influences on amplification behavior.

Inspired by Michaelis-Menten kinetics, the classical Monad model describes how growth is limited by resource consumption via a system of ordinary differential equations. An organism is assumed to reproduce at a certain theoretical maximum rate, consuming resources in the process. As resource availability decreases, the organism’s growth rate is depressed accordingly. Importantly, multiple growth-limiting substrates can be considered: the apparent growth rate is simply the product of the maximum rate with every substrate-specific reduction. The key parameters are ***r***, the maximum growth rate, ***K***_*i*_, the concentration of substrate *i* at which the apparent growth rate is half of the maximum, and ***γ*** _*i*_, the amount of organism produced per unit of substrate *i*.

In our adaptation of the Monad model, each strand of a target amplicon is treated as a separate “organism”, one which consumes primers and other substrates to “reproduce” into the complementary strand. The basic form of our model is as follows:

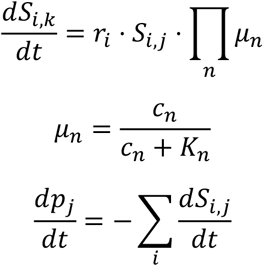

where *S*_*i,k*_ represents the concentration of the strand of amplicon *i* generated from primer *k* ; *S*_*i,j*_ the concentration of the complementary strand generated from primer *j*;*r*_*i*_ the amplification rate of amplicon *i* ; μ_*n*_ the growth limitation of reagent *n*;c_*n*_ the concentration of reagent the half-max-rate concentration of reagent ??; ??_*n*_ *n*; and the *p*_*j*_ concentration of primer *j*.

The *maximum* growth rate can be interpreted as the proportion of primer-strand complexes that are successfully converted to strand-strand duplexes via the action of the polymerase. Many factors could contribute to a sub-optimal maximum growth rate, such as amplicon length and GC content, specific sequence motifs, as well as the temperature and duration of the PCR extension phase.

By extension, the limiting effect of primer concentration on growth rate is due to the proportion of amplicon strands that bind to a primer and successfully *initiate* polymerization. This suggests that, unlike bacterial growth where a fixed substrate concentration can be assumed to limit growth to half its maximum, the concentration-at-half-max-rate *K* _*pj*_ for primers is dynamic, dependent on the thermodynamics of the primer-strand interaction.

We approximate *K* _*pj*_ to be equal to the concentration of the appropriate target strand at any given point above; because primers are typically designed to exhibit a melting temperature just above the reaction annealing temperature, we can estimate that 50% of primers will be bound to their complementary strands when both are equal in concentration at this temperature. Therefore, for primer ?? _1_ which binds to strand *S*_*N*,1_, the rate - limiting factor becomes:

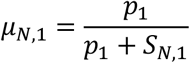

and thus (assuming no additional rate-limiting terms):

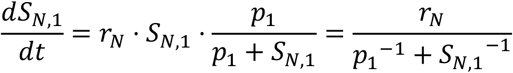

If fluorescent oligonucleotide probes *f*_*i,j*_ are used to monitor the reaction, growth of the fluorescent *signal* will be limited by the dynamics of probe-target interactions. Unlike primers, however, probes are typically designed to have a melting temperature several degrees above the PCR extension temperature. *K*_*f i,j*_ thus needs to be more carefully estimated empirically or calculated from thermodynamics. Below, *K*_*f i,j*_ is given by *κ*_*f i,j*_ *S*_*i,j*_ where ĸ is the (likely slightly > 1) multiple of the current amplicon concentration at which 50% of the initial probe population is bound to the amplicon strand. Furthermore, fluorescent yield *y*_*x*_ is needed to translate concentration of free fluorophore *x* into the observed signal. Finally, consumption of additional growth limiting reagents *R* (e.g., dNTPs) is assumed to be proportional to the number of incorporated nucleotides. Starting reagent concentration *R*_*0*_ and half-max concentration *K*_*R*_ must be estimated empirically.

### Bipartite Module

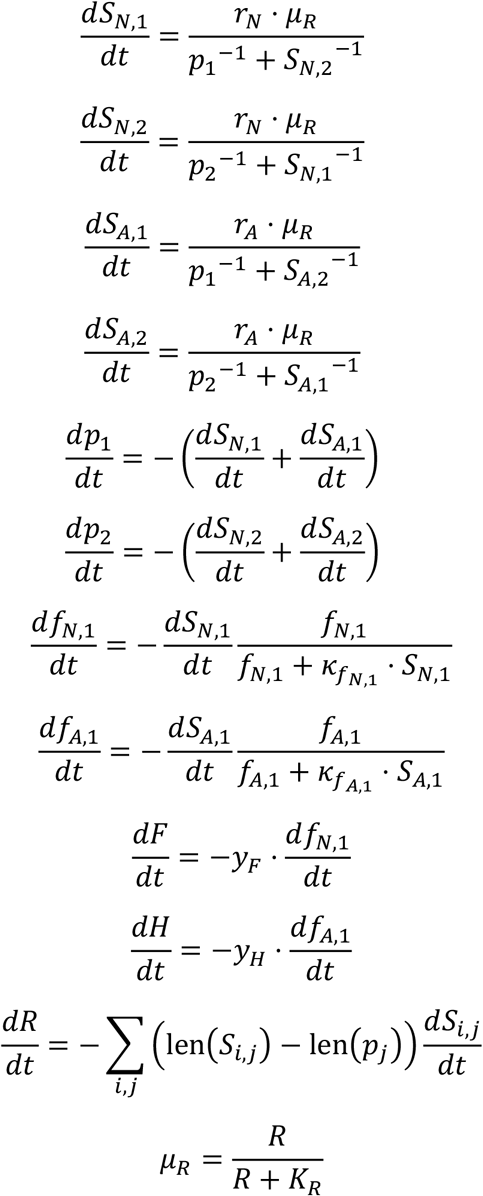

### Tripartite Module

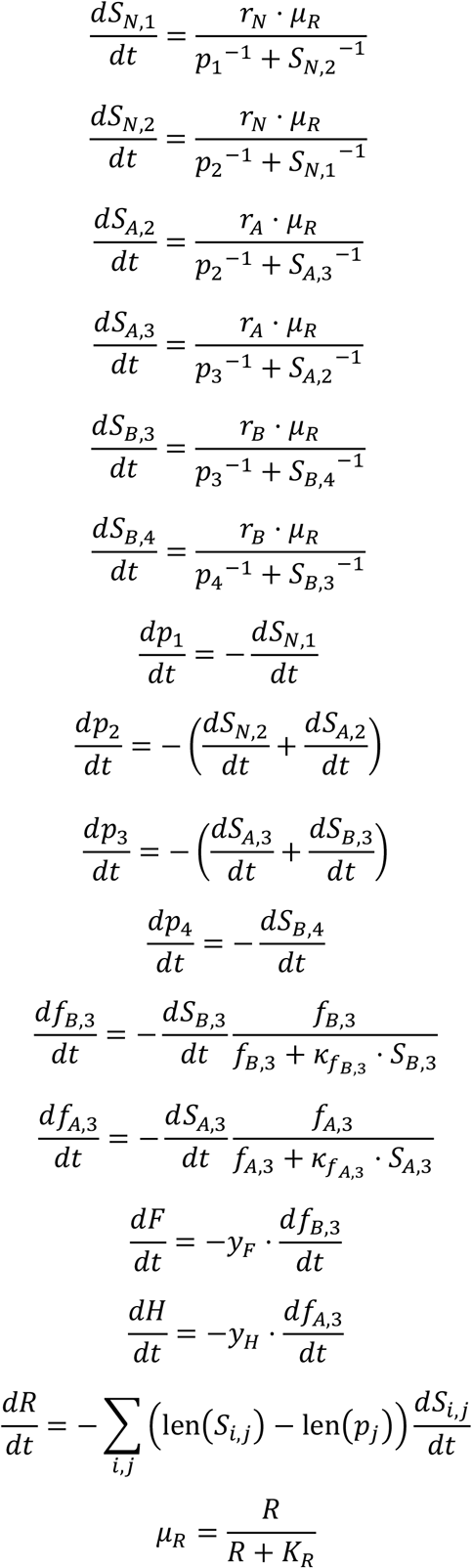

### Redundant Module (parallel)

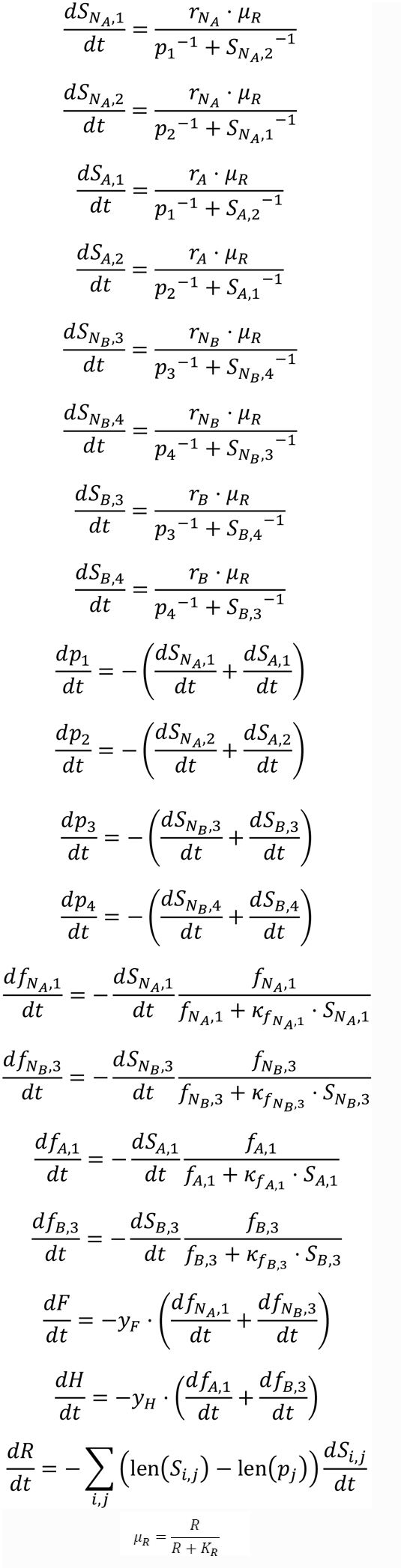

### Signal drift

An unexpected consequence of running the qPCR reactions as primer-limited (typically 1.Sx probe over primer) was the presence of significant signal drift in the post-amplification, “plateau” phase of the reaction **(Figure S6)**. While this phase is expected to remain flat, we instead observed a steady increase in the signal. This signal drift was strongly dependent on both amplicon sequence (especially GC content) and choice of probe fluorophore, quencher, and intermediate modifications. GC-rich amplicons tended to have higher degrees of drift, while xanthene dyes had higher drift than cyanine dyes and internal modifications slightly decreased drift. However, reactions monitored with EvaGreen dye displayed essentially no drift, even at 70% or greater GC content **(Figure 2B)**. Notably, the baselines for all reactions remain nearly flat for all amplicon concentrations, with signal drift only appearing *after* amplification has supposedly ceased. This signal drift is a nuisance, as it compromises the sharpness of competitive amplification response curves.

The mechanism behind this signal drift is unclear to us. The absence of drift in EvaGreen reactions rules out primer-dimers as well as amplicon self-priming. Conversely, the dependence on sequence context is evidence against a purely chemical interaction between probe labels and amplification by-products. It’s possible that the observed drift is a result of *Taq* polymerase “proofreading” of uncleaved probes. DNA polymerases are known to exhibit 3’-exonuclease activity, particularly towards unpaired bases.^50^ To speculate, perhaps the polymerase is binding to amplicon-bound probes and excising the 3’ nucleotides along with the quencher. While the synthetic targets amplified **in Figure S6** contained identical probe sequences, the rest of their inter primer sequences varied. This exact sequence context, in particular GC content, could affect secondary structure and duplex stability, and by extension both probe-amplicon binding and accessibility of the bound probe to the polymerase. Furthermore, the fluorophore and quencher modifications could affect stability of the amplicon-bound probe by anchoring the termini to different degrees. The IBRQ and BHQ2 quenchers, along with the cyanine dyes, have been shown to stabilize duplexes to a greater degree than IBFQ and the xanthene dyes.^51-53^ The observed difference between cyanine and xanthene dyes may actually be due to IBRQ/BHQ2 stabilizing the 3’ terminus of the probe, leading to decreased transitory 3’ “flap” formation and thus less polymerase proofreading than IBFQ and TAMRA. However, we fit multiple generalized linear models to these observations, and Bayesian model comparison suggested that those models which included *fluorophore* in their predictors performed better than those which included *quencher* (due to statistical confounding, models could not include both) (not shown). Further investigation is needed to illuminate the cause of this signal drift.

### Limitations of the model

In reality, *K* _*pj*_ will be a function of the current concentrations of primer *p* _*j*_ as well as all amplicons *S*_*i*,j_ utilizing *p*_*j*_. The calculation of *K*_*pj*_ could be refined by solving for the exact half-max concentration at every PCR cycle but doing so would introduce considerable algorithmic complexity. A possible middle ground could be to approximate the half-max concentration once for each primer, using the exact melting and annealing temperatures, but this is largely unnecessary owing to both the similarity between primer designs and the inherent imprecision of melting-temperature calculations. It should also be noted that we make the simplification of using deterministic continuous-time ODE solvers rather than discrete and/or stochastic solvers, which might more accurately reflect the nature of PCR but be more complex to implement.

Our model does not account for the impact of partially replicated target strands or primer dimers. In the worst case, these could act as an “unproductive” sink for primers, neither generating a signal nor producing a complementary strand with an intact primer region. However, some partial products may contain an intact probe region, and so continue to generate a signal at a linear rate. Some partial products may even be “filled in” through re-association with a complete complement and subsequent polymerase re-extension. Naturally, the impact of these pathological products will be inversely proportional to amplification rate, so low-rate amplicons will be further impaired by this mechanism.

## Extended Data Figures

**Extended Data Figure 1.**
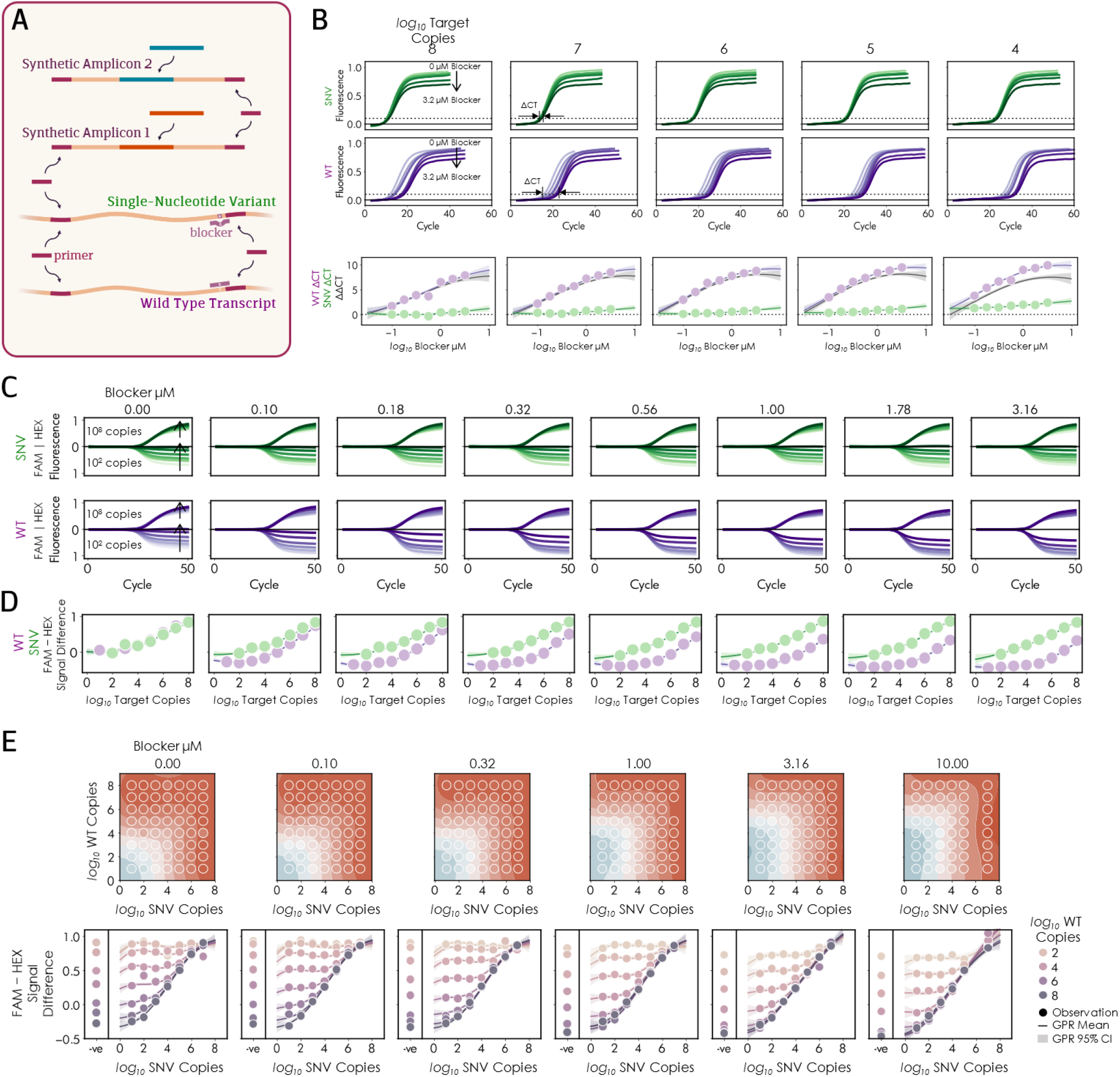
**(A)** Combining Competitive Amplification Networks with Blocker Displacement Amplification (BOA) allows detection of single nucleotide variants (SNVs) at very low variable allele frequency (VAF). Synthetic amplicons were designed to mimic the EGFR WT and the L858R SNV mutation. **(B)** The Blocker oligo delays the WT CT more than that of the SNV. **(B)** Real time fluorescence curves and **(C)** endpoint signal intensities for titrations of a “blocker” oligonucleotide designed to outcompete a primer for binding to the WT near the SNV site. High concentrations of the blocker depress the signal from the WT target greater than from the SNV. **(D)** Top and bottom rows show the same data with different visualizations: the signal response from various combinations of WT, SNV, and blocker amplified together. Observations are shown in circles while the smooth surfaces and lines are the result of Gaussian Process regression with 95% confidence intervals shown as shaded areas. In the absence of blocker, WT and SNV targets produce similar signals, as indicated by the square contours and symmetry of the regression surface (top) about the WT=SNV line. At high blocker concentrations, however, the rectangular contours indicate that the resulting signal comes entirely from the SNV target at WT concentrations less than 10^5^ copies. By inspecting the individual signal profiles of SNV detection at specific WT concentrations (bottom) we can determine the LOO as the point at which the SNV-negative reaction falls outside the 95% Cl of the respective profile. Thus, this system enables detection of mutant alleles at countable concentrations down to 0.1% VAF and up to 10^5^ copies of WT.

## Supplemental Figures

**Figure S1.**
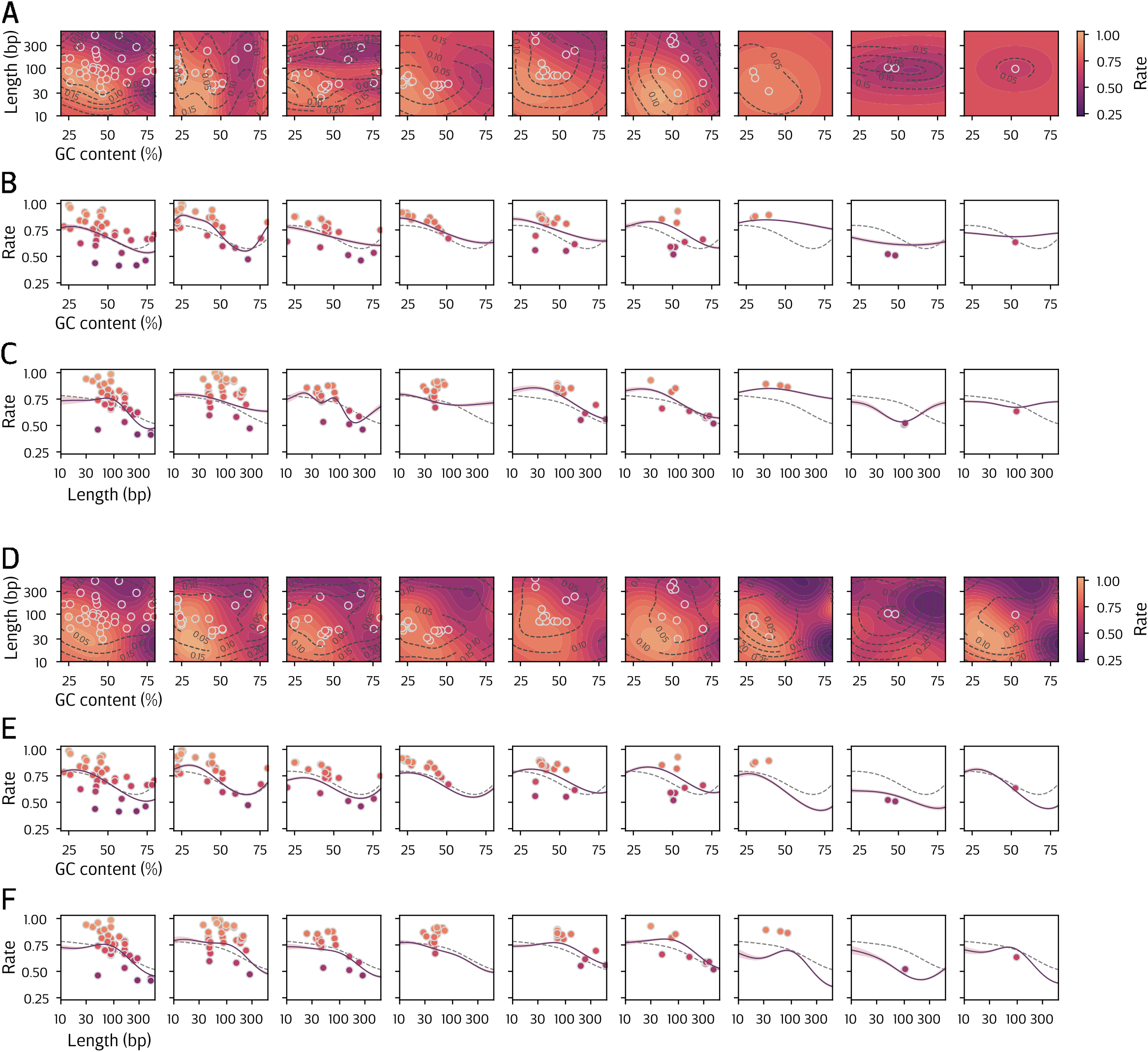
Gaussian Process regression relating amplicon length and GC content to observed amplification rate, treating each primer pair as a distinct regression surface. Each surface was fit either independently (**A-C**) or collectively through the linear model of coregionalization (**D-F**) (panel D reproduced from Figure 2D for ease of comparison). Individual observations (circles) and shown with full learned surfaces **(A, D)**, the marginal contributions (purple line) of GC content **(B, E)** and length (**C, F)** to rate inferred by each model, and the marginal contributions inferred by the global model (**Figure 2C**, dashed grey lines).

**Figure S2.**
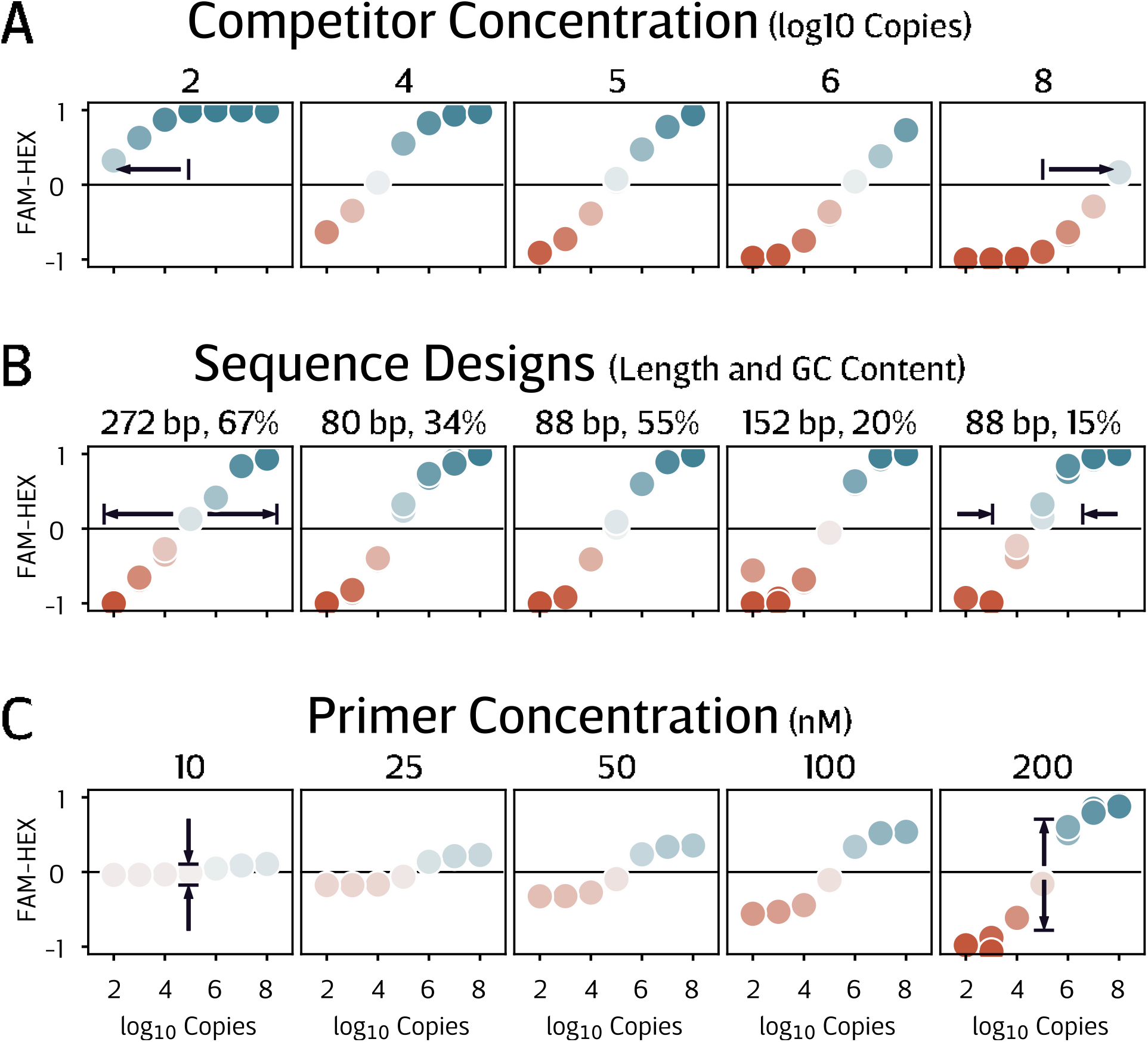
The response profile of an individual reaction module can be fine-tuned to match the desired function. **(A)** In the “direct” module shown here, the midpoint of the response can be shifted towards higher or lower target concentrations by adjusting the concentration of the competitor. **(B)** The dynamic range can be tightened or broadened through appropriate choice of the competitor length and GC content. **(C)** Finally, the magnitude of the response can be compressed or expanded via the concentration of the corresponding primers.

**Figure S3.**
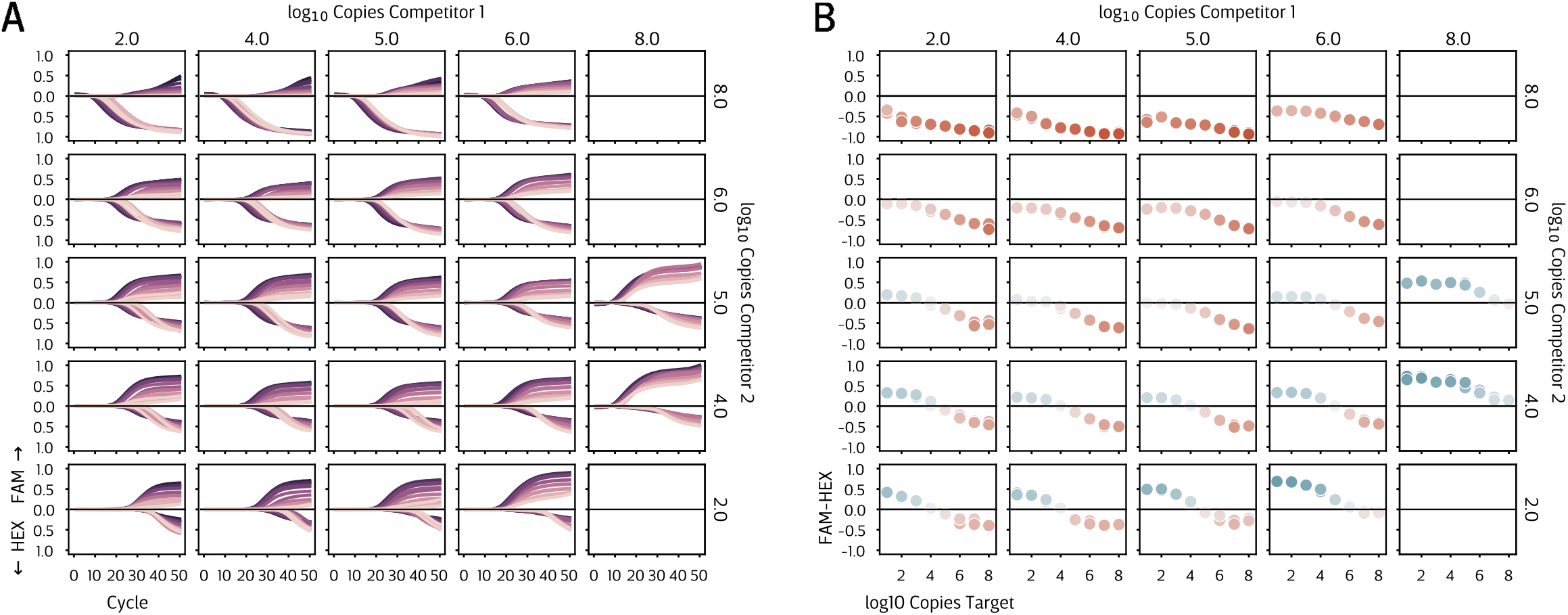
Fine-tuning of the “tripartite” module. **(A)** Real-time fluorescence curves and **(B)** endpoint response profiles for various concentrations of the two competitor sequences and the natural target. Varying the two competitor concentrations affects both the midpoint and the degree of asymmetry in the response to various natural target concentrations.

**Figure S4.**
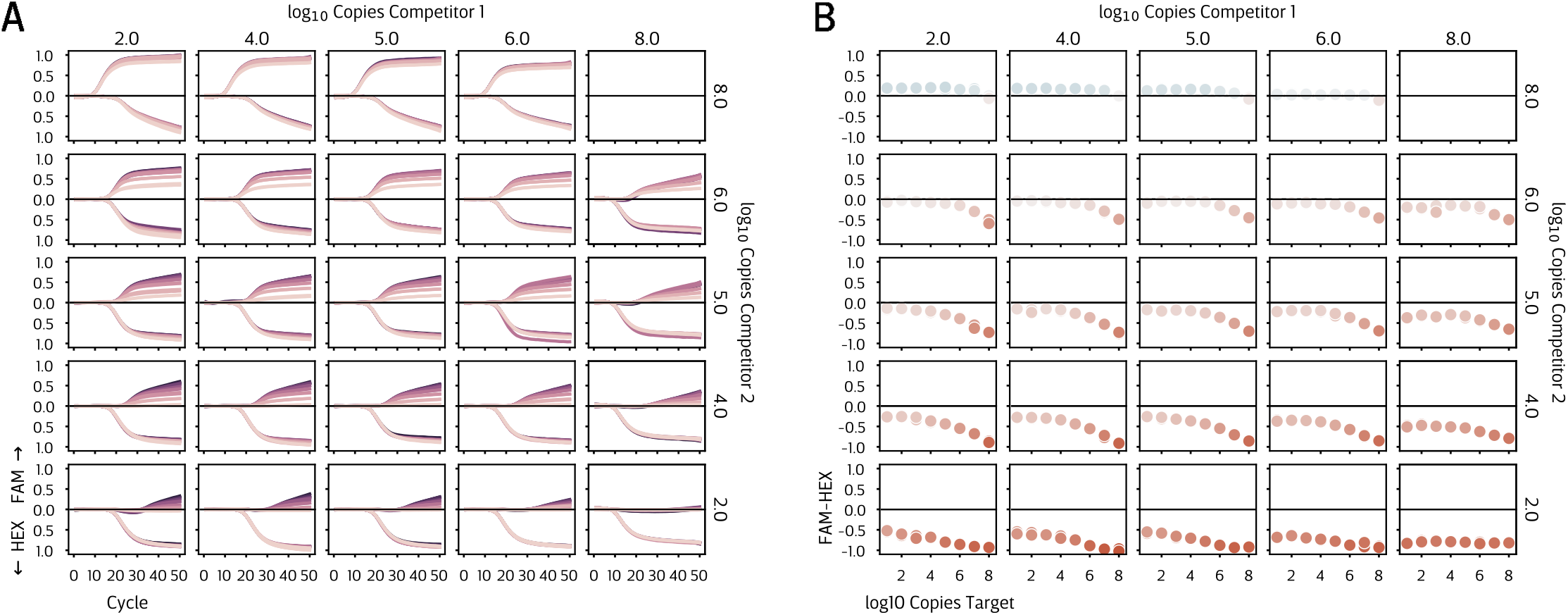
The “tripartite” module requires carefully tuned amplicon sequences to achieve desired behavior. **(A)** Real-time fluorescence curves and **(B)** endpoint response profiles for various concentrations of the two competitor sequences and the natural target. Unlike Figure 53, which used amplicon sequences exhibiting solo amplification rates near their model optimized ideals (0.82, 0.52, 0.50 for the target and competitors 1 and 2, respectively), the solo amplification rates for the sequences used here differed significantly from their ideals (0.71, 0.64, 0.65), leading to poor competitive behavior.

**Figure S5.**
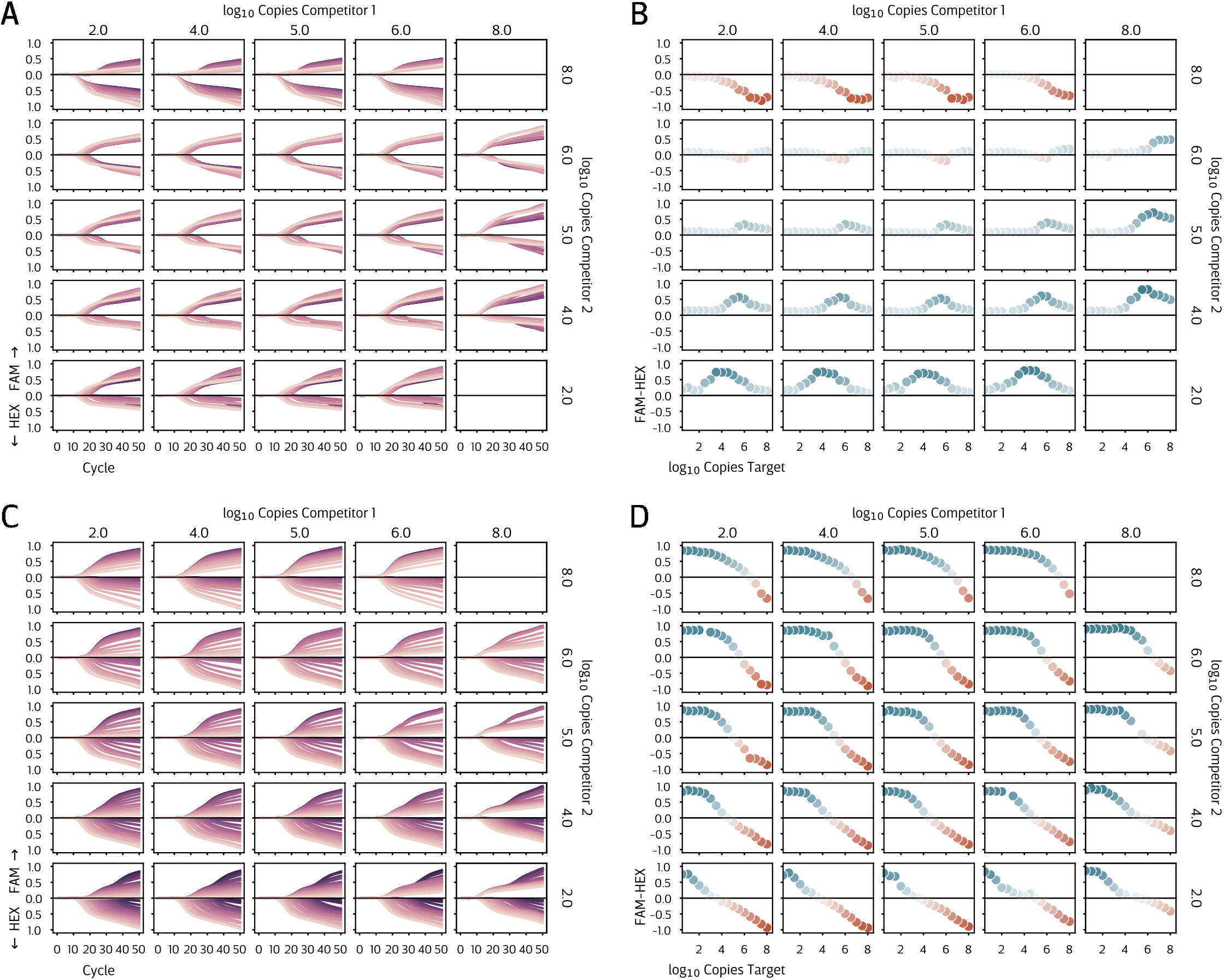
Fine-tuning of “redundant” modules, showing real-time curves **(A, C)** and endpoint signals **(B, D). (A, B) In** the “antiparallel” configuration, the two segments of the natural target bear probes with different fluorescent colors, as do the two competitors. This gives rise to a peaked response to varying concentrations of the natural target. Adjusting the concentrations of the competitors determines both the direction and location of the peak. **(C, D) In** the “parallel” configuration, the two natural target segments have probes with the same fluorescent colors. Adjusting the concentrations of the competitors can produce extended and multiphasic response profiles.

**Figure S6.**
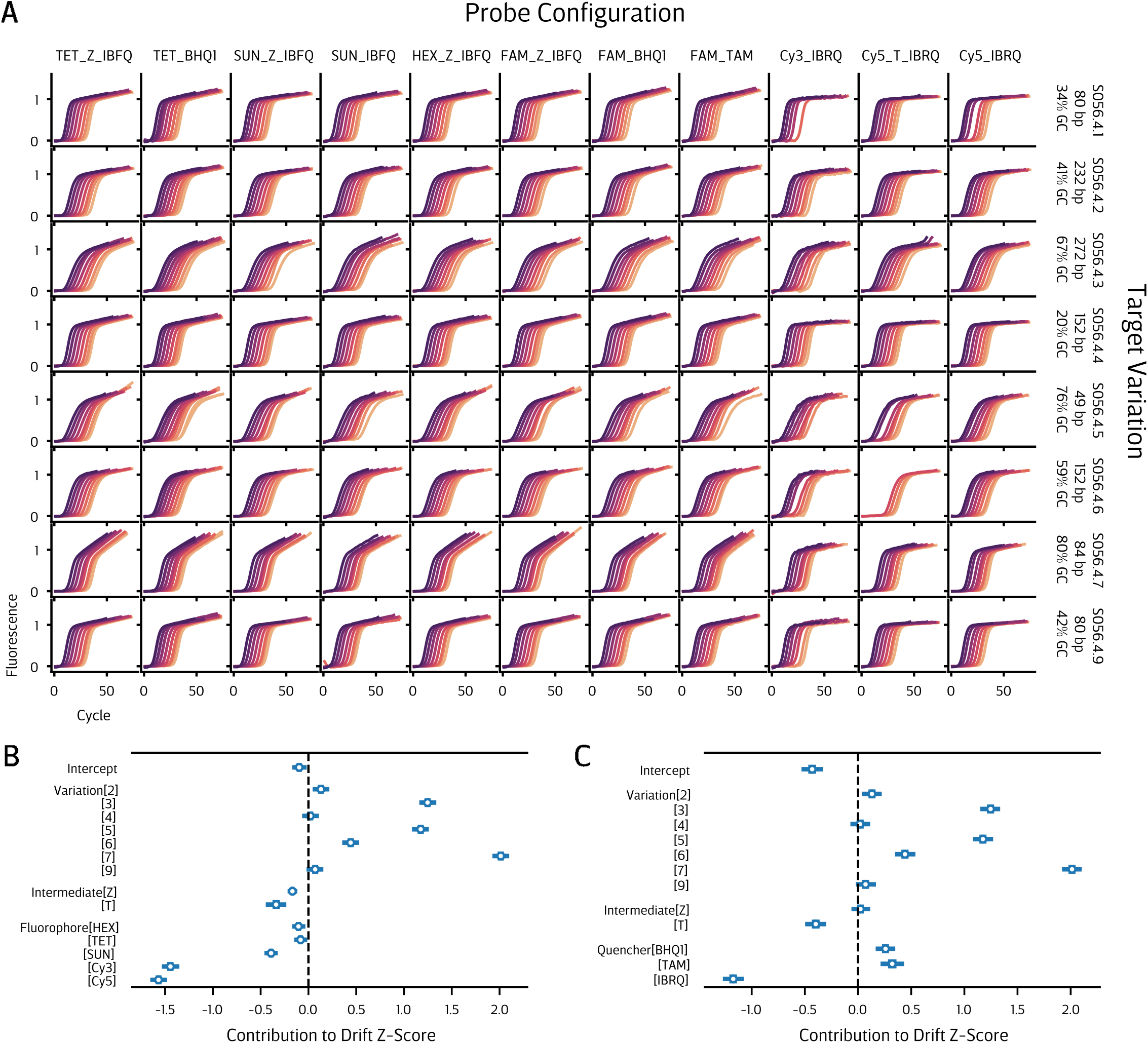
Impact of amplicon sequence and probe design on signal drift. (A) Synthetic qPCR targets were designed to utilize the same primer and probe sequences, varying in the intervening regions. These targets were amplified in primer-limited qPCR reactions with 1.5x excess of probe oligos bearing various combinations of fluorophore, intermediate, and quencher labels. The fluorescent signal is expected to level off in the post-amplification, “saturation” phase of the reaction, but instead drifts steadily upwards. **(B, C)** Generalized linear models were used to decompose the sources of this drift. The linear slope of the drift phase of each reaction was log-transformed then centered and scaled to obtain Z-scores. Then, a Bayesian linear model was fit to determine the relative impact of sequence variation and probe modifications; shown here are the resulting coefficients. Statistical confounding prevented consideration of both fluorophores and quencher simultaneously, so one model included only fluorophores as predictors **(B)** and another only quenchers **(C)**. Notably, Bayesian model comparison via PSIS-LOO indicated that the fluorophore model achieved better performance than the quencher model. The “intercept” for these models consists of the Variation 1target with a FAM-IBFQ probe. The interpretation of z-scores is such that, for example, a probe labeled with Cy5 is expected to exhibit a drift roughly 1.6 standard deviations less than one labeled with FAM for any given target, and similarly Variation 7 is expected to have a drift 2.0 standard deviations higher than Variation 1, for any given probe.

